# Disentangling the effects of Corticotrophin Releasing Factor and GABA release from the ventral bed nucleus of the stria terminalis on ethanol self-administration in mice

**DOI:** 10.1101/2023.03.02.530838

**Authors:** C.A. Gianessi, G.B. Gereau, H.L. Haun, D. Pati, T. Sides, S.L. D’Ambrosio, K. Boyt, W.P. Kelson, C.W. Hodge, T.L. Kash

## Abstract

Excessive alcohol use causes a great deal of harm and negative health outcomes. Corticotrophin releasing factor (CRF), a stress-related neuropeptide, has been implicated in binge ethanol intake and ethanol dependence. CRF containing neurons in the bed nucleus of the stria terminalis (BNST^CRF^) can control ethanol consumption. These BNST^CRF^ neurons also release GABA, raising the question, is it CRF or GABA release or both that is controlling alcohol consumption. Here, we used viral vectors to separate the effects of CRF and GABA release from BNST^CRF^ neurons on the escalation of ethanol intake in an operant self-administration paradigm in male and female mice. We found that CRF deletion in BNST neurons reduces ethanol intake in both sexes, with a stronger effect in males. For sucrose self-administration there was no effect of CRF deletion. Suppression of GABA release, via knockdown of vGAT, from BNST^CRF^ produced a transient increase in ethanol operant self-administration following in male mice, and reduced in motivation to work for sucrose on a progressive ratio schedule of reinforcement in a sex-dependent manner. Together, these results highlight how different signaling molecules from the same populations of neurons can bidirectionally control behavior. Moreover, they suggest that BNST CRF release is important for high intensity ethanol drinking that precedes dependence, whereas GABA release from these neurons may play a role in regulating motivation.

## Background

In 2014 approximately 137.9 million people in the United States were current alcohol users, of whom 43.6% were binge alcohol drinkers and 6.4% met criteria for Alcohol Use Disorder (AUD) (1). Binge drinking, which the NIAAA defines as a pattern of drinking over a short period of time that brings blood alcohol concentration levels to 0.08 g/dl, may represent an important vulnerability to the development of AUD (2,3). Escalation of drinking is a key process in the transition from casual to excessive drinking in AUD; however, the discrete mechanisms that drive this process are not fully understood. Corticotrophin releasing factor (CRF) is a stress-responsive neuropeptide that is released from the paraventricular nucleus of the hypothalamus (PVN) and from additional regions like the bed nucleus of the stria terminalis (BNST). CRF is engaged in escalated alcohol intake, theorized to be compensating for excessive reward as a major component of the pathological progression to alcohol use disorder (4, for review: 5).

Although early clinical studies of corticotrophin releasing factor receptor type 1 (CRF-R1) antagonists were unsuccessful (6), multiple independent genetic association studies have revealed a link between excessive alcohol consumption and this gene (7,8). Basic mechanistic research into how corticotrophin releasing factor (CRF) signaling is engaged over the course of escalated alcohol consumption is thus warranted, and may elucidate up or downstream regulators of CRF that may serve as successful therapeutic targets. Repeated binge drinking is one form of escalated intake that engages CRF neurons in the BNST (9–11). The majority of CRF-expressing neurons within the BNST are GABAergic (12–15). Inhibition of CRF-BNST neurons reduces binge drinking (10).

Over the past ten years, rates of AUD have increased in women by 84%, relative to a 35% increase in men (16). More research is needed into sex differences in the neurobiological response to alcohol (17). The BNST is a sexually dimorphic brain region, and differences in structure are due to organizational influences of sex hormones during early development (18,19). Additionally, sex-specific differences in CRF and GAD67 (one enzyme that synthesizes GABA) were observed within the BNST of mice (20), highlighting the importance of studying males and females. Here, we use viral vectors to selectively reduce expression of CRF or the vesicular GABA transporter (vGAT) in the ventral BNST to determine the role of CRF release or GABA release on binge ethanol intake in an operant model of self-administration in male and female mice.

## Methods and Materials

Adult male and female mice were used in the current study. Transgenic mice (CRH-Cre (21) and CRH-flox (22)) were bred inhouse. C57Bl6/J mice (Stock #: 000664, Jackson Laboratories; Bar Harbor, ME, USA) arrived at 8 weeks, and allowed to acclimate to the vivarium for at least 7 days prior to the start of the experiment. Mice were group-housed with same-sex littermates in polycarbonate cages (GM500, Techniplast; West Chester, PA, USA) on ventilation racks within a climate-controlled vivarium maintained in a 12 hr light/dark cycle (lights off at 7 am). All procedures were approved by the UNC Chapel Hill Institutional Animal Care and Use Committee and were performed in accordance with the National Institutes of Health *Guide for the Care and Use of Laboratory Animals*.

### Operant behavior

Mice were maintained at 85-90% of free-feeding body weight for the duration of the experiment by feeding 2.5-3.0 grams of standard rodent chow (Prolab Isopro RMH 3000, LabDiet; St. Louis, MO, USA) per mouse per day. Water was available *ad libitum* in the vivarium but was removed prior to behavioral testing. Mice were fed for 15 min without water available to induce thirst just prior to behavioral sessions. Before beginning instrumental training, mice were exposed to their assigned reinforcing fluid *ad libitum* (9% ethanol v/v with 2% w/v sucrose or 2% w/v sucrose alone) in their home cage for 24 hours.

Behavior was assessed during the dark cycle in 8 standard operant conditioning chambers housed within sound-attenuating cubicles (Med Associates; St. Albans, VT, USA). Each chamber was equipped with two retractable levers located on the back wall with a cue light centered above in between, with a magazine positioned in the center of the front wall. The magazine was equipped with a photobeam sensor to record entries, and a drinking trough connected to a syringe pump. Liquid reinforcers were delivered as 14 µL rewards and delivery was accompanied by a 2 second cue light. Responses during this time were counted but did not contribute to the response requirement. At the end of each session the drinking trough was checked for any residual fluid. Sessions were 45 min in duration, began with the levers extending, and ended with the levers retracting.

At the beginning of the experiment, one lever was assigned to deliver reward (referred to as “active”) and the other lever had no programmed consequence (referred to as “inactive”). Assignment of active lever was counterbalanced across mice and maintained throughout the duration of the experiment. Active lever responses were initially reinforced using a fixed ratio 1 (FR1) schedule, where each press resulted in delivery of a single reinforcer. Once individual mice earned a criterion of 10 reinforcers in a single FR1 session (1-5 days, mean 1.5 days), they were then trained on a fixed ratio 2 (FR2) schedule for three days. Mice that took longer than 2 days to achieve the performance criterion (n=6 total in all experiments) underwent remedial FR1 sessions that were extended to a duration of 2 hours. After three days of FR2 training, they were then trained on a fixed ratio 4 (FR4) schedule for 9 days.

After 9 days of FR4 training, the mice underwent progressive ratio testing to examine the motivation to work for liquid reinforcers. These progressive ratio tests were modified for use with this ethanol and sucrose solution by changing the session timeout to 90 seconds from the last earned reward, which was three times the average C57Bl6/J self-administration inter-reinforcer interval (23). First mice were tested with an arithmetic progression, where the ratios progressed 1, 1, 2, 2, 3, 3, 4, 4, 5, 5, 7, 7, 9, 9, 11, 11, 13, 13, 15, 15, 18, 18, 21, 21, 24, 24,… (24). Mice received FR4 self-administration days between progressive ratio sessions to re-establish stable responding. The second progressive ratio test followed an exponential progression where the ratios were determined using (5*e^0.2*reward^)-5, rounded up to the nearest integer (25). Breakpoint is not a continuous measure, so we report rewards earned, which is proportional to the log transformed breakpoint (23,25).

### Blood ethanol concentration

In an experiment with male and female C57Bl6/J mice, we measured blood ethanol concentration to compare this to our calculated grams per kilogram measure. Within 5 minutes of the end of the 9^th^ FR4 session mice were deeply anesthetized with isofluorane, and trunk blood was taken. Blood was centrifuged and plasma was collected. Blood ethanol concentration was measured using an Analox-AM1 alcohol analyzer (Analox Technologies, Atlanta, GA, USA).

### Stereotaxic Surgery

Adult mice were anesthetized with isoflurane (1-3%) in oxygen (1-2 l/min), and aligned on a stereotaxic frame (Kopf Instruments; Tujunga, CA, USA) while on a heated pad (Homeothermic monitoring system, Harvard Apparatus; Holliston, MA, USA). All surgeries were conducted in a sterile environment using aseptic techniques. The scalp was sterilized with 70% ethanol and betadine, then a vertical incision was made before using a drill to burr small holes in the skull directly above the injection targets. Microinjections were performed with a 1 µL Neuros Hamilton syringe (Hamilton, Reno, NV, USA) and a micro-infusion pump (Nanoject III, Drummond Scientific; Broomall, PA, USA) that infused virus at 100 nL/min. Viruses were administered bilaterally with 200 nL per side (relative to bregma: ML ±0.9 mm, AP, 0.23 mm, DV −4.75 mm), and the needle was left in place for 5 min to allow for diffusion of the virus before the needle was slowly withdrawn. Most mice were administered a single injection of buprenorphine (0.1mg/kg, s.c.) for surgical analgesia, with supplemental pain management from Tylenol water. For a total of 9 mice in behavioral studies and 4 mice in the electrophysiology experiment, the approved surgical analgesia protocol changed to ketoprofen injections (5 mg/kg, s.c.) on the day of surgery and at least two days post-surgery. Mice were allowed to recover for at least 3 weeks prior to being used for behavioral studies, and 8 weeks prior to being used for slice electrophysiology.

To selectively knockdown GABA release from CRF neurons, CRH-Cre mice were injected with either short hairpin RNAi directed at the vesicular GABA transporter, AAV8-hSyn-flex-GFP-shvGAT, or the control scrambled sequence, AAV8-hSyn-flex-GFP-shScramble (UNC Vector Core; Chapel Hill, NC, USA), as previously described (26,27). For the functional validation of this construct, each of these viruses were mixed in a 4:1 ratio with AAV8-Ef1a-double floxed-hChR2(H134R)-mCherry-WPRE-HGHpA (Addgene; Watertown, MA, USA) and injected with 300 nL per side. To selectively knock down CRF expression in the ventral BNST (28), CRH-flox mice were injected with either AAV8-hSyn-GFP or AAV8-hSyn-Cre-GFP (UNC Vector Core).

### Viral placement validation for behavioral studies

Mice were anesthetized with an overdose of the anesthetic Tribromoethanol (Avertin, 1 mL, i.p.), and transcardially perfused with chilled 0.01 M phosphate-buffered saline (PBS) followed by 4% paraformaldehyde (PFA) in PBS. Brains were extracted and post-fixed in 4% PFA for 24 hours and then stored in PBS at 4°C. 45 µm coronal sections were collected using a Leica VT1000S vibratome (Leica Biosystems; Deer Park, IL, USA) and then stored in 0.02% sodium azide (Sigma Aldrich) in PBS. The tissue was then mounted on slides and allowed to dry before cover slipping with Vecta-Shield Hardset Mounting Medium with DAPI (Vector Laboratories; Newark, CA, USA). Viral placements were imaged at 4x magnification using a Keyence BZ-X800 All-in-one Fluorescence microscope (Keyence; Itasca, IL, USA).

### Slice electrophysiology

About 8 weeks post-surgery, mice were anesthetized with isoflurane and rapidly decapitated. Coronal sections through the BNST (300 μm) were prepared as previously described. Briefly, brains were quickly extracted, and slices were made using a Leica VT 1200s vibratome (Leica Biosystems, IL, USA) in ice-cold, oxygenated sucrose solution containing in mM: 183 sucrose, 20 NaCl, 0.5 KCl, 2.5 MgCl2, 1.2 NaH2PO4, 10 glucose and 26 NaHCO3 saturated with 95 % O2/5 % CO2. Slices were incubated for at least 30 minutes in artificial cerebral spinal fluid (ACSF) maintained at 35°C that contained in mM: 124 NaCl, 4.0 KCl, 1 NaH2PO4, 1.2 MgSO4, 10 D-glucose, 2 CaCl2, and 26 NaHCO3, saturated with 95% O2/5 % CO2 before transferring to a submerged recording chamber (Warner Instruments, CT, USA) for experimental use.

Neurons were identified using infrared differential interference contrast on a Scientifica Slicescope II (Scientifica; East Sussex, UK). Fluorescent cells were visualized using a 470 nm LED. Whole-cell patch clamp recordings were performed on non-fluorescent neurons in the BNST. Identical optical stimulation parameters (5 ms of 470 nm with a power of 2.5 mW) were used to compare optically evoked currents in the BNST between the two groups. Optically evoked currents in the BNST were measured in voltage clamp using cesium methanesulfonate-based intracellular solution (Cs-Meth; in mM: 135 cesium methanesulfonate, 10 KCl, 1 MgCl2, 0.2 EGTA, 4 MgATP, 0.3 Na2GTP, 20 phosphocreatine, pH 7.3, 285–290 mOsm with 1 mg/mL QX-314) to detect light-evoked inhibitory postsynaptic currents (oIPSCs) at +10 mV. Data were sampled at 10 kHz and low pass filtered at 3 kHz. Access resistance was continuously monitored and changes greater than 20% from the initial value were excluded from data analyses.

### Statistical analysis

Data were analyzed using Prism 9 (Graphpad, San Diego, CA, USA), and SPSS 26 (IBM, Armonk, NY, USA). Data are presented in figures as the mean ± standard error of the mean. Data are presented in sex-separated graphs for clarity, sex effects were analyzed and reported in the text. Response rates and rewards earned across virus, sex, and behavioral sessions were analyzed using repeated measures GLM with a Poisson distribution, because this distribution is the most appropriate for count data. Regression coefficients were tested with Wald χ^2^ to determine if they were significantly different from zero. Significant interaction effects were analyzed pairwise among relevant conditions (sex or virus) across days with a least significant difference adjustment for multiple comparisons. Significance level was set at α= 0.05. For ethanol self-administration studies, grams per kilogram were calculated based on the volume consumed using the following formula ((0.014 L * number of rewards earned) – volume leftover in drinking trough after session) *0.09*0.789) / (body weight converted to kg). Grams per kilogram of ethanol consumed, and blood ethanol concentration were analyzed using the normal distribution since these are continuous measures. Blood ethanol concentration and calculated grams per kilogram were correlated with a simple linear regression. Percentage of responsive neurons in viral conditions was compared using Fisher’s exact test. Amplitude and latency of oIPSCs were analyzed with unpaired t tests between viral conditions, with Welch’s correction applied for unequal variances when appropriate.

## Results

### Operant Binge Ethanol Self-Administration

High levels of ethanol consumption are achieved in our operant self-administration model for 9% (v/v) ethanol with 2% sucrose in food restricted, post-prandial mice during the dark cycle. The level of ethanol and sucrose included in the ethanol solution is about equivalent to a Moscato wine. Male and female C57BL/6J mice responded on the active lever to self-administer ethanol (Figure 1A). Females tended to respond at higher rates than males did across days, with a significantly higher response rate on day 9 of FR4 (Main effect of sex: Wald χ^2^(1)=3.4, p=0.06; Main effect of day: Wald χ^2^(8)=253.1, p<0.0001; Sex-by-day interaction: Wald χ^2^(8)=32.9, p<0.0001; day 9 p<0.05). Females have a smaller body mass than males do, which translated to significantly higher calculated grams per kilogram consumption of ethanol across days of FR4 (Figure 1B.

**Figure 1:**
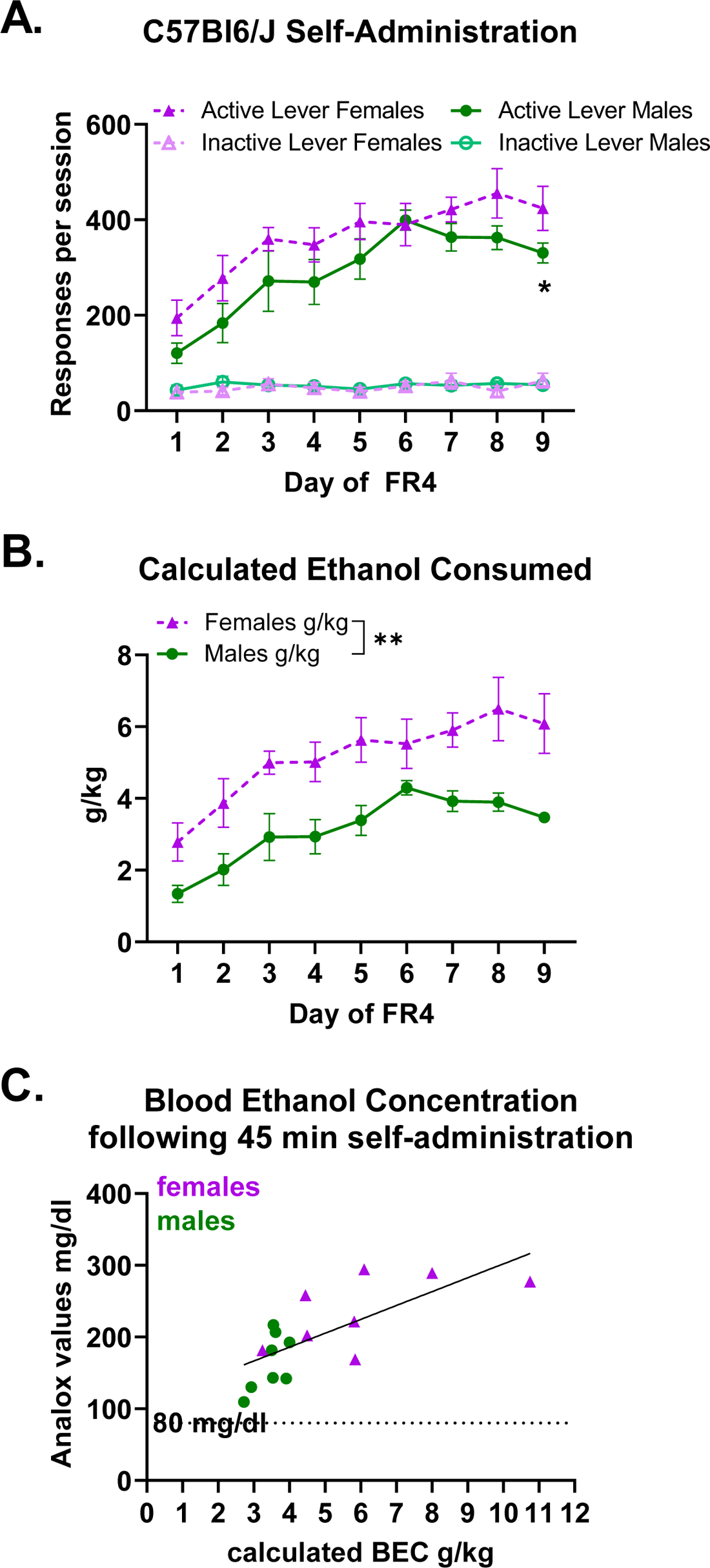
Operant Binge Ethanol Self-Administration model a. Male (n=8) and female (n=8) C57Bl6/J mice learn to respond on the active lever for ethanol reinforcers. Females tended to respond on the active lever more than males did across days, with the effect being significant on day 9. Main effect of sex: Wald χ2 (1) = 3.4, p=0.06; Main effect of day: Wald χ2 (8)= 253.1, p<0.0001; Sex-by-day interaction: Wald χ2 (8) = 32.9, p<0.0001. Day 1 p=0.07, Day 2 p= 0.1, day 3 p=0.2, day 4 p=0.2, day 5 p=0.14, day 6 p=0.8, day 7 p=0.1, day 8 p=0.08, day 9 p<0.05 b. Female C57Bl6/J mice consumed more ethanol on a grams per kilogram basis. 2way ANOVA g/kg: Main effect of Sex F(1,14) = 12.9, p=0.003; Main effect of Day F(8) = 16.1, p<0.0001; Sex-by-day interaction F(8,112) = 0.8, p = 0.6. Note that the standard error of the mean for Day 9 FR4 males is too small to be depicted (Mean 3.47± 0.16 SEM) c. On the ninth day of FR4, all mice had blood ethanol concentrations above 80 mg/dl, which is the threshold for binge intoxication. The blood ethanol concentration values significantly correlated with the calculated grams per kilogram consumed, R2 =0.5233, p=0.0015

2way ANOVA g/kg: Main effect of Sex F(1,14)=12.9, p=0.003; Main effect of Day F(8)=16.1, p<0.0001; Sex-by-day interaction F(8,112)=0.8, p=0.6). Blood ethanol concentrations were measured immediately following the 9^th^ FR4 self-administration session (Figure 1C). All of the mice achieved above the binge intoxication threshold, and the ethanol concentrations in the blood correlated with the calculated ethanol consumed (R^2^=0.52, p=0.0015).

### Ethanol self-administration ventral BNST CRF Knockdown

Knocking down expression of CRF peptide within the ventral bed nucleus of the stria terminalis led to reductions in ethanol self-administration. This effect was more pronounced in males: when active lever responses across days are tested for sex and virus effects there is a significant sex-by-virus interaction (Wald χ^2^(1)=9.0, p=0.003) and a sex-by-virus-by-day interaction (Wald χ^2^(8)=15.3, p=0.05). When examining the GFP-expressing control mice, was no significant main effect of sex, or sex-by-day interaction, which indicates that the sex effects are driven by the CRF knockdown groups. When examining the CRF knockdown mice, there was a significant main effect of sex (Wald χ^2^(1)=11.6, p< 0.0001) and a significant sex-by-day interaction (Wald χ^2^ (8)=43.0, p<0.0001). Male CRF knockdown mice had significantly lower active lever responding than female CRF knockdown mice on all days except day 4. Male CRF knockdown mice self-administered less ethanol than male GFP controls (Figure 2A. Main effect of virus Wald χ^2^(1)=18.9, p<0.0001). There was a trend for a reduction in active lever responding in female CRF Knockdown mice (Figure 2B. Main effect of virus: Wald χ^2^(1)=3.3, p=0.07).

**Figure 2:**
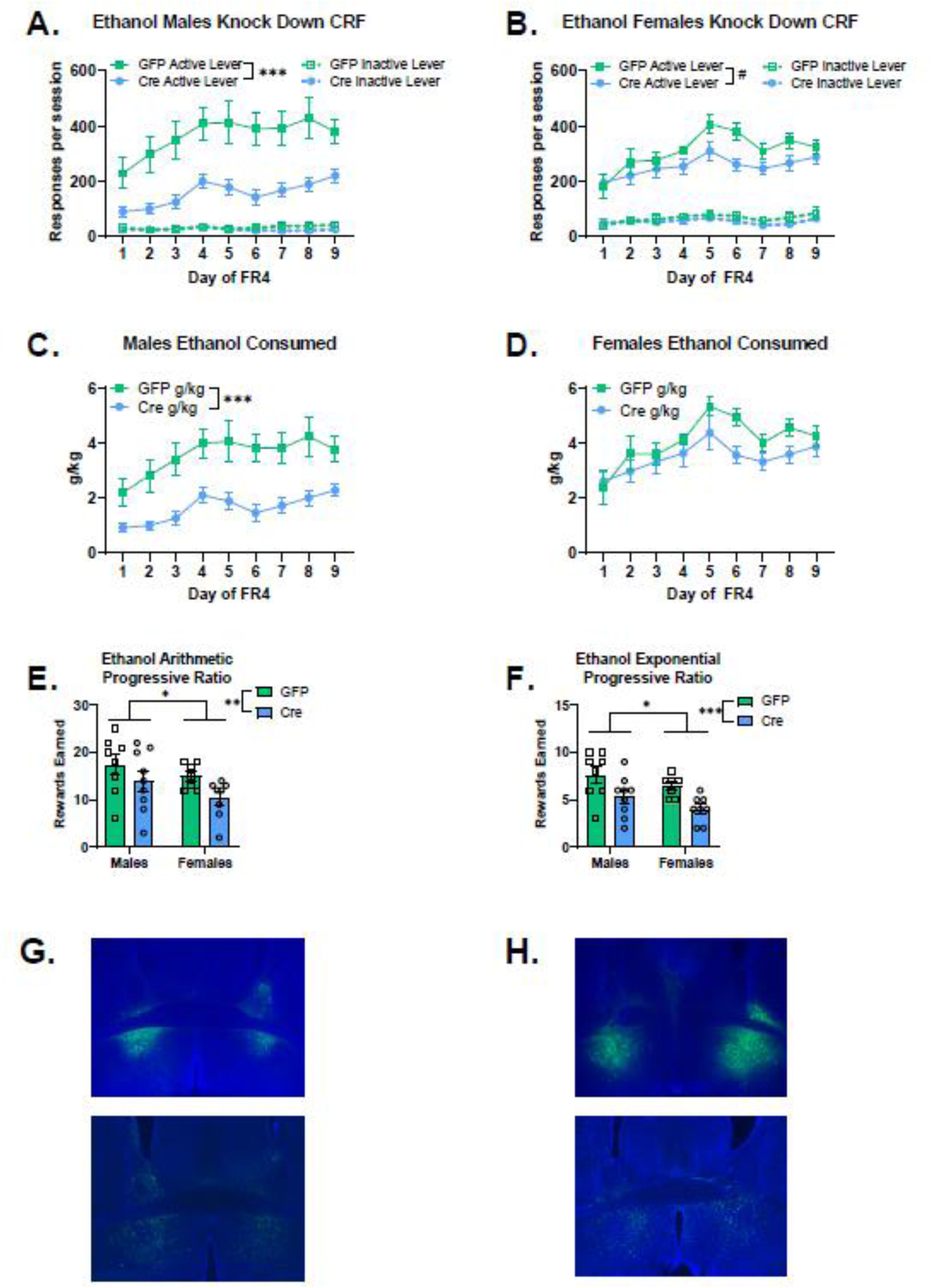
CRF Knockdown in ventral BNST reduced ethanol self-administration, with a larger effect in males a. Males CRF knockdown mice (n=9) self-administered less ethanol than male GFP controls (n=8). Main effect of virus Wald χ^2^ (1) = 18.9, p<0.0001 b. Female CRF knockdown mice (n=8) had a trend for reduced active lever responding on an FR4 schedule compared to GFP controls (n=7). Main effect of virus: Wald χ^2^ (1) = 3.3, p = 0.07 c. Male CRF knockdown mice consumed less ethanol than male GFP controls. Main effect of virus: Wald χ^2^ (1) = 12.8, p<0.0001, DayFR4 x Virus: Wald χ^2^ (8) = 72.5, p<0.0001, all days’ p≤ 0.01 d. Female CRF knockdown mice had a trend for reduced ethanol consumption compared to female GFP controls. Day-by-virus interaction Wald χ^2^ (8) = 14.7, p=0.066, lower g/kg values on day 6 (p=0.001) and day 8 (p=0.02). e. CRF knockdown reduced motivation to work for ethanol reinforcers on an arithmetic progressive ratio schedule. Female mice earned fewer rewards on this progressive ratio test than males did. Rewards earned: Main effect of virus: Wald χ^2^ (1) = 6.4, p=0.01, Main effect of sex: Wald χ^2^ (1) = 3.8, p=0.05 f. CRF knockdown reduced motivation to work for ethanol reinforcers on an exponential progressive ratio schedule. Female mice earned fewer rewards on this progressive ratio test than males did. Main effect of virus: Wald χ^2^ (1) = 15.6, p<0.0001, Main effect of sex: Wald χ^2^ (1) = 4.7, p = 0.03 g. Representative images of viral placement in male mice expressing the control GFP virus (top) and the Cre expressing virus (bottom). h. Representative images of viral placement in female mice expressing the control GFP virus (top) and the Cre expressing virus (bottom).

Knocking down expression of the CRF peptide also reduced the consumption of ethanol as measured by g/kg, with a similar pattern to the active lever responding. When assessing g/kg across days of FR4 by sex and virus, there is a significant main effect of sex (Wald χ^2^(1)=13.1, p<0.0001), and sex-by-virus interaction (Wald χ^2^(1)=4.1, p=0.04). Examining the role for sex in the GFP-expressing control mice revealed a significant sex-by-day interaction (Wald χ^2^(8)=18.6, p=0.017), where female GFP mice had significantly higher g/kg on day 6 (p<0.05) compared to male GFP mice, reflecting the tendency for female mice to consume higher levels of ethanol than males. Examining the role for sex in the Cre-expressing control mice revealed a significant main effect of sex (Wald χ^2^(1)=28.5, p<0.0001) and sex-by-day interaction (Wald χ^2^(8)=28.5, p<0.0001), where male CRF knockdown mice consumed less ethanol than female CRF knockdown mice did (all days’ p<0.01). Male CRF knockdown mice consumed less ethanol than male GFP controls (Figure 2C. Main effect of virus: Wald χ^2^(1)=12.8, p<0.0001, virus-by-day interaction: Wald χ^2^(8)=72.5, p<0.0001, all days’ p≤ 0.01). There was a tendency for female CRF knockdown mice to consume less ethanol than female GFP controls, reflected in a trend for a day-by-virus interaction (Figure 2D. Wald χ^2^(8)=14.7, p=0.066), and significantly lower g/kg values on day 6 (p=0.001) and day 8 (p=0.02).

Knocking down CRF peptide expression in the ventral BNST reduced motivation to work for ethanol reinforcers in the progressive ratio tests. Rewards earned on the arithmetic progressive ratio test (Figure 2E) were significantly lower for CRF knockdown animals (Main effect of virus: Wald χ^2^(1)=6.4, p=0.01) and also significantly lower for females (Main effect of sex: Wald χ^2^(1)=3.8, p=0.05). The same pattern of results was observed in the exponential progressive ratio test (Figure 2F. Main effect of virus: Wald χ^2^(1)=15.6, p<0.0001, Main effect of sex: Wald χ^2^(1)=4.7, p=0.03).

### Sucrose self-administration ventral BNST CRF Knockdown

Sucrose self-administration was not affected by knockdown of CRF peptide in the BNST. When examining active lever responses across days by sex and virus, there was no significant main effect of virus (Wald χ^2^(1)=0.012, p=0.9) or sex-by-virus interaction (Wald χ^2^(1)=1.7, p=0.2). There was a significant main effect of sex (Wald χ^2^(1)=20.2, p<0.0001), and sex-by-virus-by-day interaction (Wald χ^2^(8)=15.7, p<0.05), due to higher levels of active lever responses in females compared to males. When examining GFP-expressing control mice, females responded more for sucrose than GFP-expressing males did (main effect of sex: Wald χ^2^(1)=14.5, p<0.0001; sex-by-day interaction: Wald χ^2^(8)=62.3, p<0.0001). Females with CRF knockdown responded more for sucrose than CRF knockdown males (Main effect of sex: Wald χ^2^(1)=6.1, p=0.014; sex-by-day interaction: Wald χ^2^(8)=55.4, p<0.0001). Males escalated their responding across days of FR4, whereas females had elevated FR4 responding earlier.

There was no significant effect of CRF knockdown among active lever responses in males self-administering sucrose (Figure 3A. Main effect of virus: Wald χ^2^(1)=0.64, p=0.4). Also, there was no difference in active lever responding for sucrose between virus conditions in females (Figure 3B. Main effect of virus: Wald χ^2^(1)=1.7, p=0.2).

**Figure 3:**
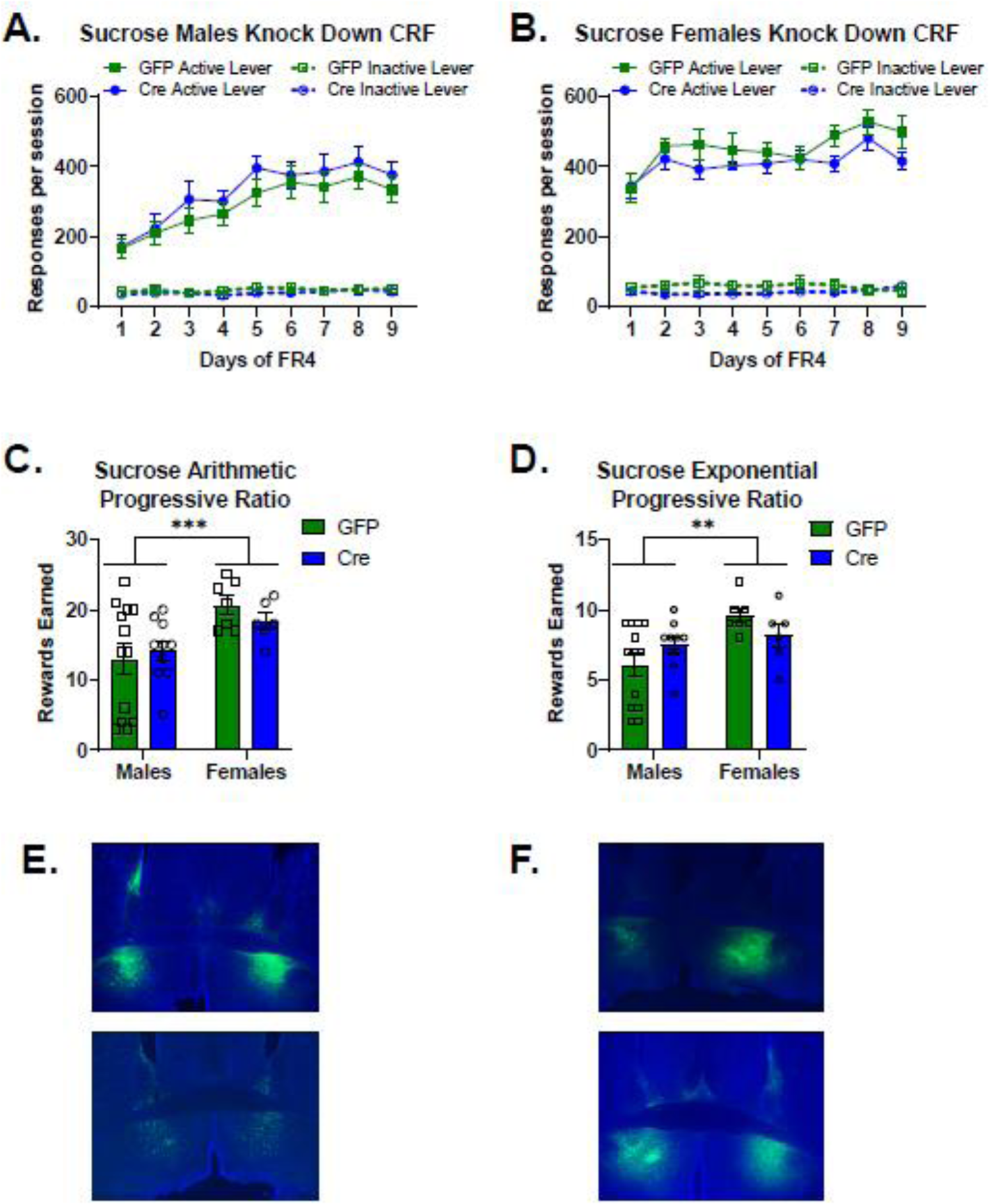
CRF knockdown in the ventral BNST did not alter sucrose self-administration a. There was no significant effect of CRF knockdown among active lever responses in males self-administering sucrose (Male GFP n=13, Cre n=10). Main effect of virus: Wald χ^2^ (1) = 0.64, p = 0.444 b. There was no significant difference between virus conditions in females in active lever responding for sucrose (Female GFP n=7, Cre n=6). Main effect of virus: Wald χ^2^ (1) = 1.7, p=0.2 c. CRF knockdown did not affect rewards earned in the arithmetic progressive ratio test. Females had higher motivation to work for sucrose reinforcers on this schedule. Main effect of sex: Wald χ^2^ (1) = 12.9, p<0.0001 d. CRF knockdown did not alter rewards earned in the exponential progressive ratio test. Main effect of virus: Wald χ^2^ (1)= 0.86, p=0.8. Females earned more rewards on this schedule than males did. Main effect of sex: Wald χ^2^ (1)= 9.6, p=0.002. There was a significant sex-by-virus interaction: Wald χ^2^ (1) = 4.5, p=0.03, however, there were no significant effects of virus for males (p=0.1) or for females (p=0.1) on rewards earned in the exponential progressive ratio test. e. Representative images of viral placement in male mice expressing the control GFP virus (top) and the Cre expressing virus (bottom). f. Representative images of viral placement in female mice expressing the control GFP virus (top) and the Cre expressing virus (bottom).

Knocking down CRF peptide expression in the ventral BNST did not alter motivation to work for sucrose reinforcers in the progressive ratio tests. In the arithmetic progressive ratio test, there was a significant main effect of sex (Figure 3C. Main effect of sex: Wald χ^2^(1)=12.9, p<0.0001) reflecting the higher level of rewards earned in females. In the exponential progressive ratio test, there was a significant main effect of sex (Figure 3D. Main effect of sex: Wald χ^2^(1)=9.6, p=0.002, sex-by-virus interaction: Wald χ^2^(1)=4.5, p=0.03). There were no significant effects of virus for males (p=0.1) or for females (p=0.1) on rewards earned in the exponential progressive ratio test.

### GABA knockdown functional validation

Using short hairpin RNA interference directed at the vesicular GABA transporter (referred to as shvGAT), we were able to selectively reduce GABA release from CRF neurons in the BNST (26,27). We injected a mixture of channelrhodopsin and either the shvGAT virus or the scrambled sequence control virus into the BNST of CRF-Cre mice to measure optically evoked inhibitory currents. Neurons in the BNST are densely inter-connected (29,30), so whole-cell patch clamp recordings from non-fluorescent were able to detect reductions in GABA release from CRF neurons with GABA knockdown. There were no significant differences in the percentage of responsive neurons recorded from scramble controls (Figure 4A) or shvGAT-expressing mice (Figure 4B). There was a significant reduction in oIPSC amplitude in shvGAT-expressing mice compared to scramble controls (Figure 4C. t(7.6)=3.6, p=0.007), indicating that GABA release was reduced by the viral manipulation. There was no difference in oIPSC latency between viral conditions (Figure 4D. t(13)=1.5, p=0.15).

**Figure 4:**
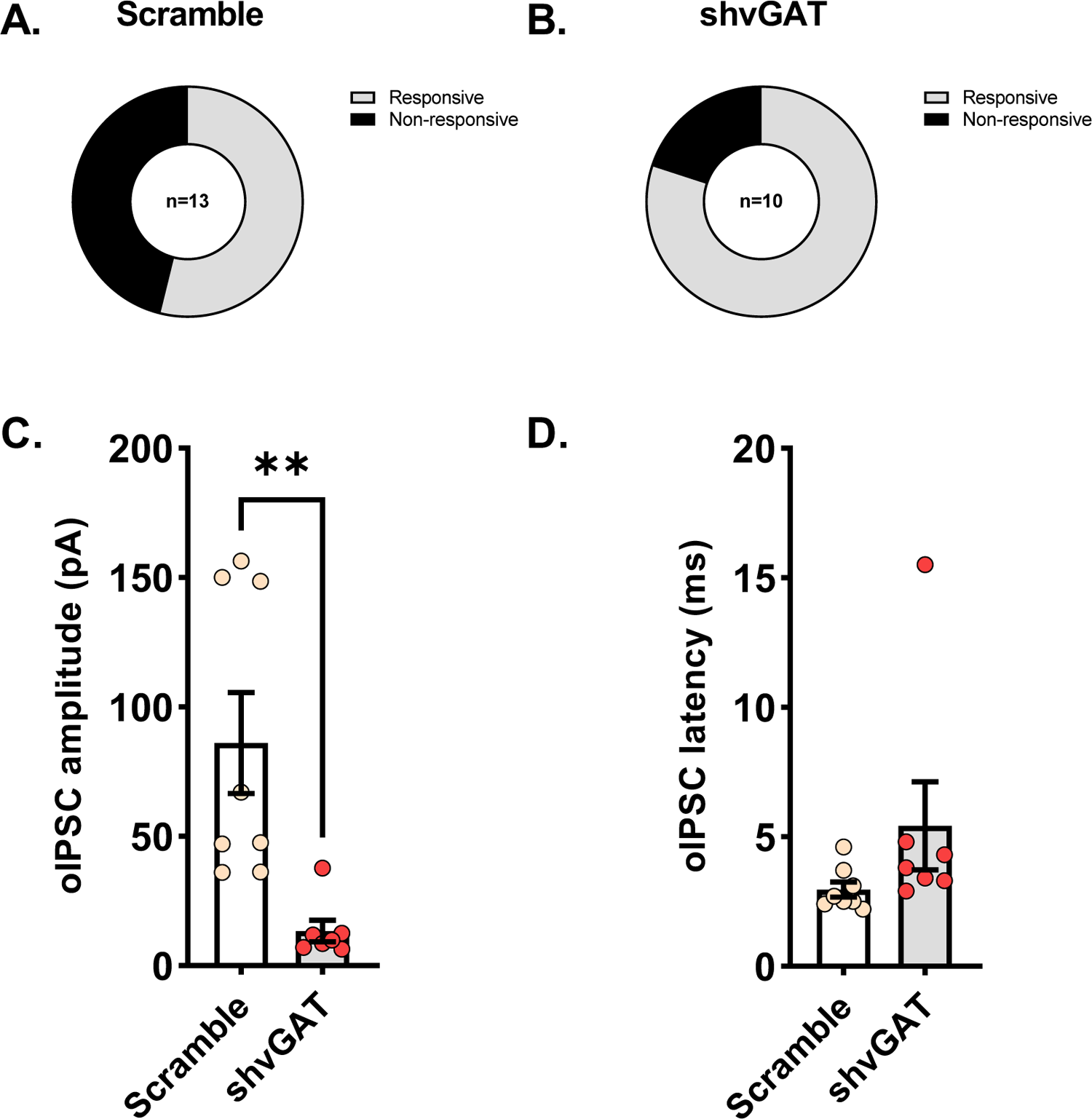
Short hairpin RNA interference for the vesicular GABA transporter (shvGAT) reduces GABA release from ventral BNST CRF-expressing neurons a. Percentage of light-responsive non-fluorescent neurons in the BNST of male mice (n=2) injected with the control virus with a scrambled sequence b. Percentage of light-responsive non-fluorescent neurons in the BNST of male mice (n=2) injected with the shvGAT virus. The percentages of responsive neurons were not different between virus conditions. Fisher’s exact test p=0.4 c. The shvGAT virus reduced optically evoked inhibitory post-synaptic current amplitude compared to the scramble controls. t(7.6) = 3.6, p=0.007 d. There were no differences in latency for optically evoked inhibitory post synaptic currents between viral conditions t(13) = 1.5, p=0.15

### Ethanol self-administration following BNST^CRF^ neuron vGAT knockdown

Knocking down GABA release from BNST^CRF^ neurons led to a transient increase in ethanol self-administration in males. When analyzing active lever responding across days for effects of sex and virus, there was a significant main effect of virus (Wald χ^2^(1)=22.0, p<0.0001), a significant sex-by-virus interaction (Wald χ^2^(1)=9.0, p=0.003), and a significant sex-by-virus-by-day interaction (Wald χ^2^(8)=15.3, p=0.05). Examining the effect of sex within the scramble virus control animals, we observed a significant sex-by-day interaction (Wald χ^2^(8)=22.0, p=0.005), reflecting the tendency for female scramble mice to respond more for ethanol between day 2 and 5 of FR4 (day 2 p=0.06, day 3 p=0.08, day 4 p=0.05, day 5 p=0.08). Within the shvGAT-expressing animals, there was a significant effect of sex (Wald χ^2^(1)=4.1, p=0.04), and a significant sex-by-day interaction (Wald χ^2^(8)=17.6, p=0.02), reflecting the tendency for male shvGAT mice to respond more for ethanol than female shvGAT mice (day 1 p=0.06, day 2 p=0.08, day 7 p=0.01, day 8 p=0.03). Male mice with reduced GABA release from BNST^CRF^ neurons self-administered significantly more ethanol compared to control males (Figure 5A. Main effect of virus: Wald χ^2^(1)=3.8, p=0.05; virus-by-day interaction: Wald χ^2^(8)=48.9, p<0.0001), and with significantly more active lever responses on the first three days of FR4 (day 1 p=0.016, Day 2 p=0.018, day 3 p=0.024, day 4 p=0.06). Reducing GABA release from BNST^CRF^ neurons did not alter female ethanol active lever responding on an FR4 schedule of reinforcement (Figure 5B. Main effect of virus: Wald χ^2^(1)=0.63, p=0.4, virus-by-day interaction: Wald χ^2^(8)=11.6, p=0.2).

**Figure 5:**
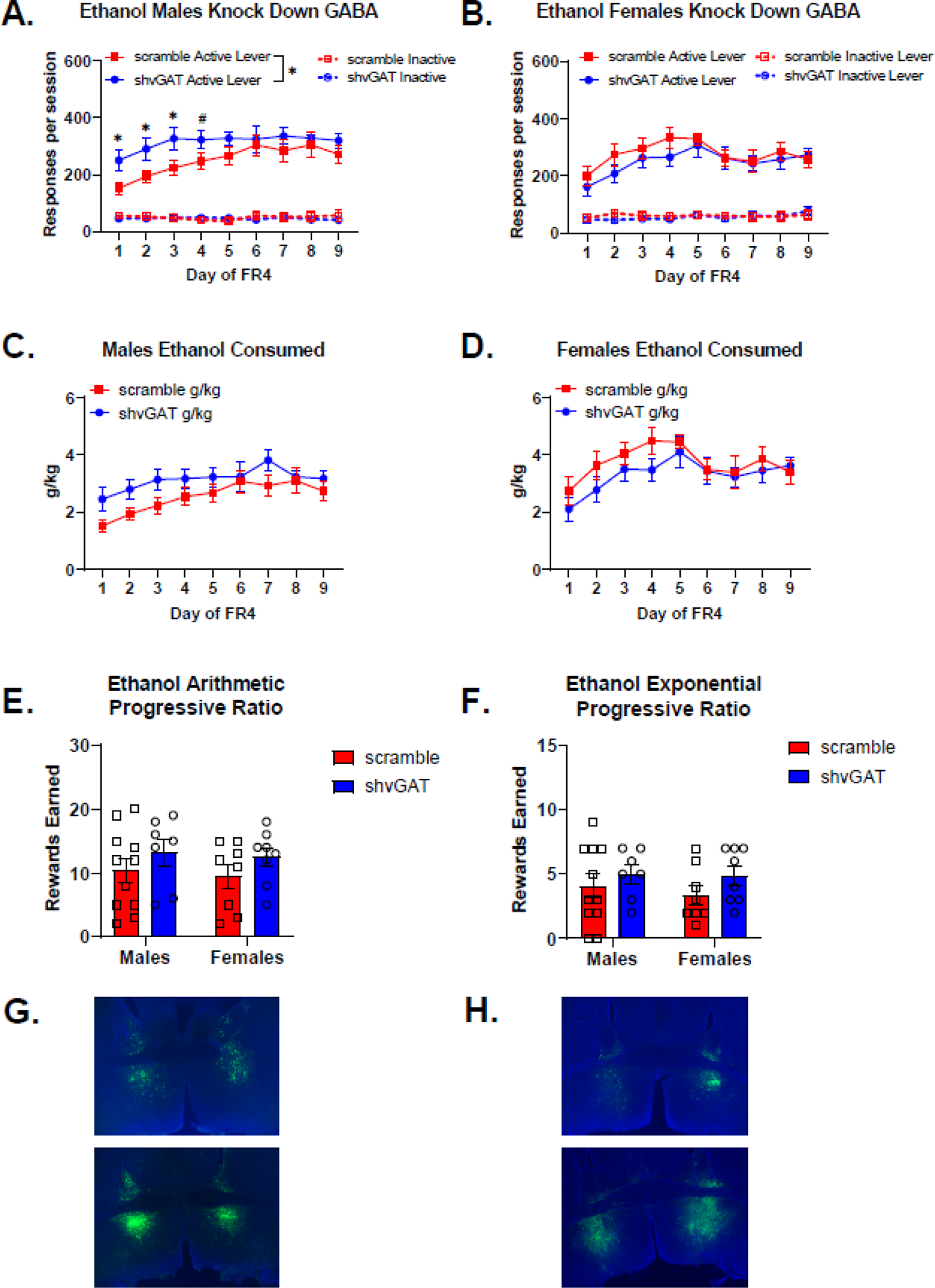
GABA knockdown in ventral BNST CRF-expressing neurons transiently increases ethanol self-administration in male mice a. Male mice with reduced GABA release from BNST-CRF neurons (male shvGAT n=7) self-administered significantly more ethanol compared to control males (male scramble n=11). Main effect of virus: Wald χ^2^ (1) = 3.8, p=0.05; virus-by-day interaction: Wald χ^2^ (8) = 48.9, p<0.0001, and with significantly more active lever responses on the first three days of FR4 (day 1 p=0.016, Day 2 p=0.018, day 3 p=0.024, day 4 p=0.06). b. Reducing GABA release from BNST-CRF neurons did not alter female ethanol active lever responding on an FR4 schedule of reinforcement (Females scramble n=8, shvGAT n=8). Main effect of virus: Wald χ^2^ (1) = 0.63, p=0.4, virus-by-day interaction: Wald χ^2^ (8) = 11.6, p=0.2. c. Male mice with GABA knockdown in BNST CRF neurons tended to consume more ethanol, but this did not reach significance. Wald χ^2^ (1) = 3.4, p=0.07. d. Female mice with GABA knockdown in BNST CRF neurons consumed similar amounts of ethanol as control females. Main effect of virus: Wald χ^2^ (1) = 0.8, p=0.4, virus-by-day interaction: Wald χ^2^ (8)= 11.4, p=0.2. e. There were no significant effects of virus or sex for the arithmetic progressive ratio test. Main effect of virus: Wald χ^2^ (1) = 2.7, p=0.1, Main effect of sex: Wald χ^2^ (1) = 0.25, p=0.6, sex-by-virus interaction: Wald χ^2^ (1) = 0.012, p=0.9. f. There were no significant effects of virus or sex for the exponential progressive ratio test. Main effect of virus: Wald χ^2^ (1) = 2.5, p=0.1, Main effect of sex: Wald χ^2^ (1) = 0.4, p=0.5, Sex x Virus: Wald χ^2^ (1) = 0.2, p=0.6. g. Representative images of viral placement in male mice expressing the control scramble virus (top) and the shvGAT expressing virus (bottom). h. Representative images of viral placement in female mice expressing the control scramble virus (top) and the shvGAT expressing virus (bottom).

Knocking down GABA release from BNST^CRF^ neurons had less of significant effect on g/kg consumed of ethanol. When examining a role for sex and virus across days of FR4, there was a significant main effect of sex (Wald χ^2^(1)=5.7, p=0.02) and a trend for a sex-by-virus interaction (Wald χ^2^(1)=3.3, p=0.07). Within scramble control mice, there was a significant effect of sex (Wald χ^2^(1)=7.8, p=0.005) and a significant sex-by-day interaction (Wald χ^2^(8)=53.9, p<0.0001). Female scramble control mice consumed significantly more ethanol than male scramble controls during the first 5 days of FR4 self-administration (day 1 p=0.02, day 2 p=0.001, day 3 p<0.0001, day 4 p<0.0001, day 5 p<0.0001). While there was a significant sex-by-day interaction for g/kg of ethanol consumed in shvGAT-expressing mice (Wald χ^2^(8)=17.2, p=0.03), none of the post hoc tests reached significance. Male mice with GABA knockdown in BNST CRF neurons tended to consume more ethanol, but this did not reach significance (Figure 5C. Wald χ^2^(1)=3.4, p=0.07). Female mice with GABA knockdown in BNST CRF neurons consumed similar amounts of ethanol as control females (Figure 5D). In our progressive ratio tests there were no significant effects of virus or sex for either the arithmetic (Figure 5E) or the exponential (Figure 5F) versions.

### Sucrose self-administration following BNST^CRF^ neuron vGAT knockdown

Reducing GABA release from BNST^CRF^ neurons had various impacts on sucrose self-administration. For active lever responding for sucrose, when analyzing the effect of virus and sex across days, there was a significant main effect of sex (Wald χ^2^(1)=5.8, p=0.02), and a significant sex-by-virus-by-day interaction (Wald χ^2^(8)=16.2, p=0.04). Analyzing active lever responding for scramble control mice, there was a significant sex-by-day interaction (Wald χ^2^(8)=24.1, p=0.002), where females tended to respond more on the active lever than males did, reaching significance on day 8 (p=0.02) and day 9 (p=0.001). When examining active lever responding for shvGAT-expressing animals, there was a significant main effect of sex (Wald χ^2^(1)=4.8, p=0.03), and a significant sex-by-day interaction (Wald χ^2^(8)=30.0, p<0.0001), with females responding more on the active lever than males (day 2 p=0.03, day 3 p=0.04, day 7 p=0.02, day 9 p=0.005). Within male mice (Figure 6A), there was a significant virus-by-day interaction (Wald χ^2^(8)=34.3, p<0.0001), which is driven by the strong main effect of day (Wald χ^2^(8)=146.0, p<0.0001), as none of the individual days were significantly different between GABA knockdown males and controls. Within female mice (Figure 6B), there was a similar pattern of results, with a significant main effect of day (Wald χ^2^(8)=171.1, p<0.0001) and virus-by-day interaction (Wald χ^2^(8)=20.1, p=0.01), yet none of the individual days were significantly different between GABA knockdown females and controls.

**Figure 6:**
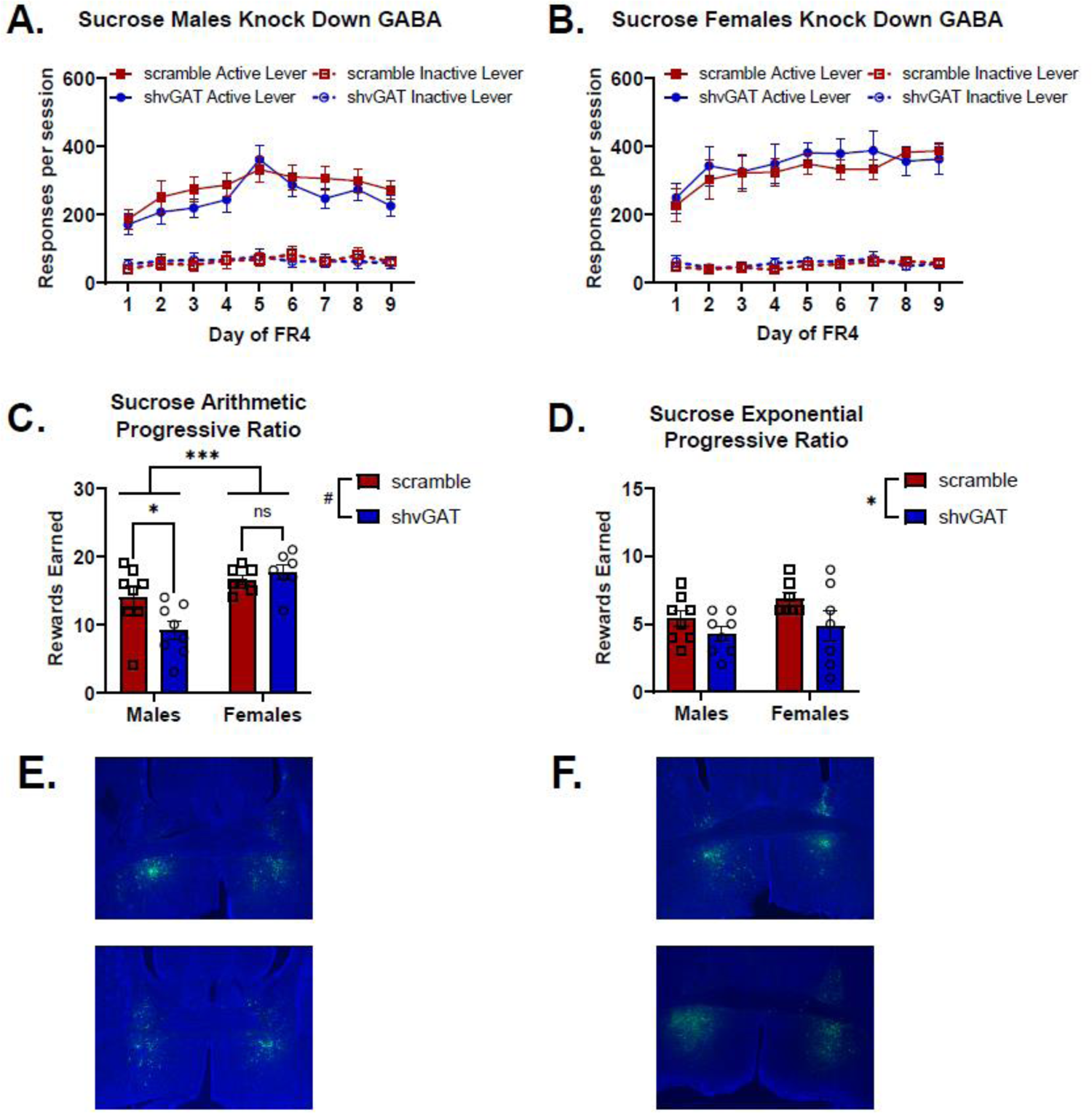
GABA knockdown in CRF-expressing BNST neurons reduced motivation to work for sucrose in a sex-dependent way a. BNST-CRF GABA knockdown did not alter sucrose self-administration in male mice (male scramble n=8, shvGAT n=8). Main effect of virus: Wald χ^2^ (1) = 0.7, p=0.4, Main effect of day: Wald χ^2^ (8) = 146.0, p<0.0001, virus-by-day interaction: Wald χ^2^ (8) = 34.3, p<0.0001. Comparing male shvGAT mice to male scramble mice across days: day 1 p=0.7, day 2 p=0.4, day 3 p=0.2, day 4 p=0.4, day 5 p=0.6, day 6 p=0.6, day 7 p=0.2, day 8 p=0.6, day 9 p=0.2 b. BNST-CRF GABA knockdown did not alter sucrose self-administration in female mice (female scramble n=7, shvGAT n=7). Main effect of virus: Wald χ^2^ (1) = 0.2, p=0.7, Main effect of day: Wald χ^2^ (8) = 171.1, p<0.0001, virus-by-day interaction: Wald χ^2^ (8) = 20.1, p=0.01. Comparing female shvGAT mice to female scramble mice across days: day 1 p=0.7, day 2 p=0.6, day 3 p=0.96, day 4 p=0.7, day 5 p=0.4, day 6 p=0.3, day 7 p=0.4, day 8 p=0.5, day 9 p=0.6 c. BNST-CRF GABA knockdown reduced motivation to work for sucrose reinforcement on an arithmetic progressive ratio schedule of reinforcement in male mice. Main effect of sex Wald χ^2^ (1) = 19.1, p<0.0001, main effect of virus Wald χ^2^ (1) = 3.6, p=0.06, sex-by-virus interaction Wald χ^2^ (1) = 6.7, p=0.009. Effect of virus in males p=0.015, effect of virus in females p=0.3. d. BNST-CRF GABA knockdown reduced motivation for both sexes to work for sucrose on an exponential progressive ratio schedule of reinforcement. Main effect of sex: Wald χ^2^ (1) = 1.9, p=0.2, Main effect of virus: Wald χ^2^ (1) = 4.5, p=0.04, sex-by-virus interaction: Wald χ^2^ (1) = 0.16, p=0.69 e. Representative images of viral placement in male mice expressing the control scramble virus (top) and the shvGAT expressing virus (bottom). f. Representative images of viral placement in female mice expressing the control scramble virus (top) and the shvGAT expressing virus (bottom).

**Figure 7:**
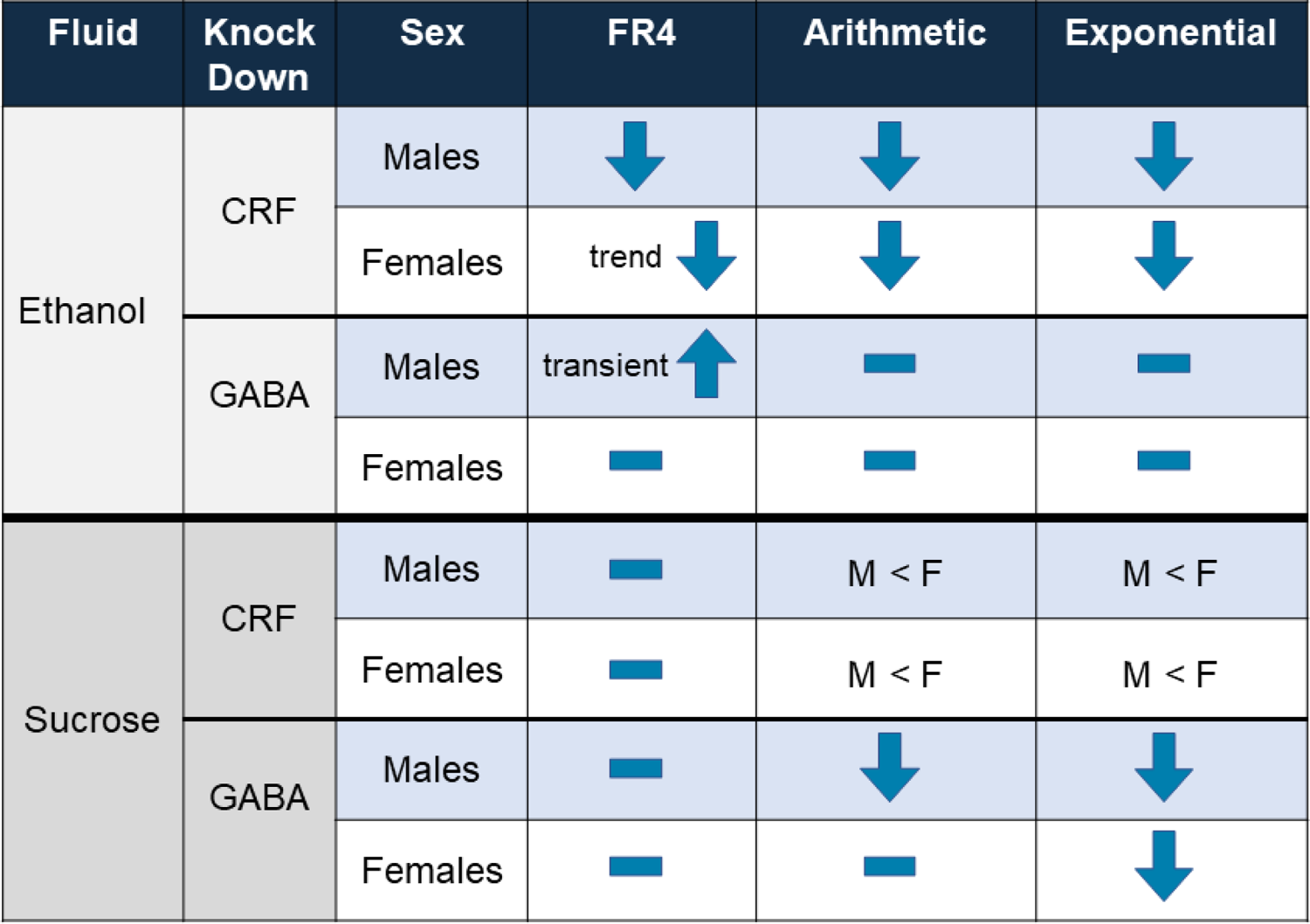
Summary of behavioral results

Knocking down GABA release from BNST^CRF^ neurons had a larger effect on males than females on motivation to work for sucrose reinforcement. In the arithmetic progressive ratio test (Figure 6C) there was a significant main effect of sex (Wald χ^2^(1)=19.1, p<0.0001), a trend for a main effect of virus (Wald χ^2^(1)=3.6, p=0.06), and a significant sex-by-virus interaction (Wald χ^2^(1)=6.7, p=0.009). GABA knockdown in BNST CRF neurons in males reduced motivation to work for sucrose on an arithmetic progressive ratio test (p=0.015), but did not alter motivation to work for sucrose in females on this test (p=0.3). In the exponential progressive ratio test, there was a significant main effect of virus (Wald χ^2^(1)=4.5, p=0.04), indicating that GABA knockdown reduced motivation to work for sucrose for both males and females.

## Discussion

In the present study, we found that deletion of CRF from the BNST led to a reduction of operant ethanol self-administration in male mice and reduced motivation to consume alcohol in both male and female mice. This appeared selective for ethanol, as we did not see similar effects of CRF knockdown when examining operant sucrose self-administration in parallel behavior-matched controls. Suppression of vGAT from BNST^CRF^ neurons transiently increased ethanol responses in male, but not female mice. Additionally, suppression of vGAT from BNST^CRF^ neurons reduced motivation to work for sucrose reinforcers, with females showing a reduction in motivation only on the exponential progressive ratio schedule. Together, these findings suggest that CRF release from BNST neurons supports high intensity drinking that precedes ethanol dependence. Suppressing GABA release from BNST^CRF^ neurons may produce a state of anhedonia, which may underlie the transient increase in ethanol self-administration in males due to stronger negative reinforcement from the ethanol, as well as the reduction in motivation for sucrose reinforcers. This study supports a model in which different compounds released from the same population of neurons can bidirectionally influence behavior. In support of this model, a recent report demonstrated opposing functions in anxiety-like behavior for CRF and GABA released from central amygdala CRF-expressing neurons in rats (27). This is an important consideration to take into account when designing and interpreting experiments that drive activation and inhibition of neuronal populations.

Previous work in our lab found that CRF-containing neurons in the BNST appeared to be important for binge-like ethanol consumption. Specifically, we found that activating a Gi-coupled DREADD led to reduced alcohol consumption, in both male and female mice (9,10). These BNST CRF neurons are GABAergic as well, raising the question was it the CRF or the GABA, or both that was driving increased alcohol intake. In male mice, there was a clear model that emerged, with CRF driving alcohol responding and motivation, and GABA playing an opposing role on alcohol responding. In females, it was less clear. Deletion of CRF led to a reduction in motivation, but minimal impact on responding. This is interesting, as in a recent study we found that there were sex differences in the physiological properties of BNST CRF neurons, with the neurons from female mice being more excitable with greater synaptic excitation (31).

Reducing GABA release from BNST CRF neurons in male mice produced a transient increase in self-administration of ethanol, but did not alter motivation to work for ethanol on our progressive ratio tests. It is possible that if we had tested for motivation differences earlier in FR4 training, we may have seen an increase. It is possible that there is some form of compensatory mechanism at play to adjust ethanol self-administration behavior following GABA release reductions, which seems more likely to happen with fast neurotransmitters like GABA compared to neuromodulators like CRF. Taken together, these data sets support a model in which there may be different mechanisms and circuits engaged to support high level alcohol consumption. These sex-differences are important to consider translationally, as there have been negative results reported for CRFR1 antagonists clinical trials for AUD that have focused on female patient populations (32).

Recent work in rats utilizing an operant two-choice task where both options gave reinforcement, but one option gave reinforcement paired with optogenetic laser stimulation demonstrated that pairing stimulation of BNST-CRF neurons with sucrose pellet reinforcement (33) led to avoidance of that operant response, whereas pairing stimulation of BNST-CRF neurons with cocaine reinforcement did not alter choice between operant responses (34). Previous work using fiber photometry recordings has shown that food consumption increases (35), and cocaine administration decreases (36), bulk neural activity in the BNST. Combined with our results, this might indicate that GABA release from BNST-CRF neurons may lessen the motivational value of sucrose reinforcers. For cocaine reinforcers, the role for decreased BNST-CRF neural activity is not related to motivation, so optogenetic stimulation does not alter cocaine-motivated behavior.

### Potential caveats of our model

Food restriction impacts the mouse behavior in this study, as does the inclusion of sweetener. Notably, in humans this is the case also. Food insecurity risk is associated with moderate and severe alcohol use disorder diagnosis in young adults (37). According to the CDC, about half of American adults drink sugar-sweetened beverages on a given day (38), and about 1 in 5 American adults is dieting, which often involves some form of food restriction (39). Indeed, the term “drunkorexia” was coined 15 years ago to describe patterns of behavior involving food restriction and binge drinking of alcohol in humans that is associated with higher risk for alcohol-related problems (40). The model used herein, rather than studying the impact of these factors in isolation to determine their individual effect, is using them as a model for common human behaviors and social determinants of health that represent important risk factors for adverse outcomes. Other operant studies for ethanol (25,41), food (42–45), and other rewards including cocaine (34,46,47) utilize food restriction as a means to enhance motivation. It is possible that enhanced motivation through food restriction may produce a ceiling effect for measuring increases in operant responding.

## Conclusion

These data demonstrate that the same genetically-defined neural population within a brain region can influence motivated behavior in opposing ways depending on which neurotransmitter is released. Additionally, neural circuits underlying motivated behavior are sexually dimorphic.

## Acknowledgements

This research was supported by grants from the National Institutes of Health’s (NIH) National Institute of Alcohol Abuse and Alcoholism (NIAAA): U01AA020911-06, U24AA025475-01, R01AA019454 (TK), R01AA028782 (CWH), P60AA011605 (TK and CWH), and T32AA007573-22 (CAG).

